# The complex quorum sensing circuitry of *Burkholderia thailandensis* is both hierarchically and homeostatically organized

**DOI:** 10.1101/128819

**Authors:** Servane Le Guillouzer, Marie-Christine Groleau, Eric Déziel

## Abstract

The genome of the bacterium *Burkholderia thailandensis* encodes for three complete LuxI/LuxR-type quorum sensing (QS) systems: BtaI1/BtaR1 (QS-1), BtaI2/BtaR2 (QS-2), and BtaI3/BtaR3 (QS-3). The LuxR-type transcriptional regulators BtaR1, BtaR2, and BtaR3 modulate the expression of target genes in association with various *N*-acyl-_L_-homoserine lactones (AHLs) as signaling molecules produced by the LuxI-type synthases BtaI1, BtaI2, and BtaI3. We have systematically dissected the complex QS circuitry of *B. thailandensis* strain E264. Direct quantification of octanoyl-homoserine lactone (C_8_-HSL), *N*-3-hydroxy-decanoyl-homoserine lactone (3OHC_10_-HSL), and *N*-3-hydroxy-octanoyl-homoserine lactone (3OHC_8_-HSL), the primary AHLs produced by this bacterium, was performed in the wild-type strain and in QS deletion mutants. This was compared to the expression of *btaI*1, *btaI*2, and *btaI*3 using chromosomal mini-CTX-*lux* transcriptional reporters. Furthermore, transcription of *btaR*1, *btaR*2, and *btaR*3 was monitored by quantitative reverse-transcription PCR (qRT-PCR). We observed that C_8_-HSL, 3OHC_10_-HSL, and 3OHC_8_-HSL are differentially produced over time during bacterial growth and correlate with the *btaI*1, *btaI*2, and *btaI*3 genes expression profiles, revealing a sequential activation of the corresponding QS systems. Moreover, transcription of the *btaR*1, *btaR*2, and *btaR*3 genes is modulated by AHLs, showing that their regulation depend on themselves, and on other systems. We conclude that the three QS systems in *B. thailandensis* are interdependent, suggesting that they cooperate dynamically and function in a concerted manner in modulating the expression of QS target genes through a sequential regulatory network.

**Importance:** Quorum sensing (QS) is a widespread bacterial communication system coordinating the expression of specific genes in a cell density-dependent manner and allowing bacteria to synchronize their activities and to function as multicellular communities. QS plays a crucial role in bacterial pathogenicity by regulating the expression of a wide spectrum of virulence/survival factors and is essential to environmental adaptation. The results presented here demonstrate that the multiple QS systems coexisting in the bacterium *Burkholderia thailandensis*, considered as the avirulent version of the human pathogen *Burkholderia pseudomallei* and thus commonly used as an alternative study model, are hierarchically and homeostatically organized. We found these QS systems finely integrated into a complex regulatory network, including transcriptional and post-transcriptional interactions, and further incorporating growth stages and temporal expression. These results provide a unique, comprehensive illustration of a sophisticated QS network and will contribute to a better comprehension of the regulatory mechanisms that can be involved in the expression of QS-controlled genes, in particular those associated with the establishment of host-pathogen interactions and acclimatization to the environment.

## Introduction

Quorum sensing (QS) is a global regulatory mechanism of gene expression depending on bacterial density (1). Gram-negative bacteria typically possess homologues of the LuxI/LuxR system initially characterized in the bioluminescent marine bacterium *Vibrio fischeri* (2). The signaling molecules *N*-acyl-L-homoserine lactones (AHLs) produced by the LuxI-type synthases accumulate in the environment throughout bacterial growth, providing information on cell density. These AHLs activate the LuxR-type transcriptional regulators that modulate the expression of QS target genes, which usually contain a *lux*-box sequence in their promoter region. These genes include a *luxI* homologue encoding a LuxI-type synthase generally located in close vicinity of a *luxR* homologue that codes for a LuxR-type transcriptional regulator, resulting in a typical self-inducing loop of AHLs (3).

Species belonging to the *Burkholderia* genus generally carry a unique AHL-based QS system referred as the CepI/CepR QS system (4). The CepI synthase is responsible for the production of octanoyl-HSL (C_8_-HSL), whereas the CepR transcriptional regulator modulates the expression of QS target genes in association with C_8_-HSL, including the *cepI* gene (4). Additionally, *cepR* expression can be auto-regulated (5, 6). Multiple QS circuitries were also described in several *Burkholderia* spp., such as the members of the *Btpm* group that consists of the non-pathogenic soil saprophyte *Burkholderia thailandensis* and the closely-related pathogens *Burkholderia pseudomallei* and *Burkholderia mallei* responsible for melioidosis and glanders, respectively (7-9). QS was reported to be involved in the regulation of several virulence factors in *B. pseudomallei* and to be essential to its pathogenicity (10). Designated as the avirulent version of *B. pseudomallei* (11), *B. thailandensis* is commonly used as a surrogate model for the study of *B. pseudomallei*, who is considered a potential bioterrorism agent and whose manipulation is consequently restricted to biosafety level 3 (BSL3) labs. The members of the *Bptm* group contain homologous LuxI/LuxR QS systems that are involved in the biosynthesis of various AHLs (12-16). In *B. thailandensis*, the LuxI/LuxR QS systems are referred as the BtaI1/BtaR1 (QS-1), BtaI2/BtaR2 (QS-2), and BtaI3/BtaR3 (QS-3) QS systems. The QS-1, QS-2, and QS-3 systems are also found in *B. pseudomallei*, whereas the QS-2 system is absent in *B. mallei* (17). These species also possess additional orphan *luxR* homologs, namely *btaR*4 (*malR*) and *btaR*5 in *B. thailandensis* E264 (7-9, 18).

The QS-1 system is composed of the *btaI*1 and *btaR*1 genes that codes for the BtaI1 synthase and the BtaR1 transcriptional regulator, respectively. BtaI1 is responsible for the production of C_8_-HSL (12, 14, 16), and transcription of *btaI*1 is positively modulated by BtaR1 (19). The BtaI2 synthase and the BtaR2 transcriptional regulator encoded by the *btaI*2 and *btaR*2 genes, respectively, constitute the QS-2 system. BtaR2 directly activates expression of *btaI*2 involved in both 3OHC_10_-HSL and 3OHC_8_-HSL biosynthesis (15). The QS-3 system comprises the *btaI*3 gene encoding the BtaI3 synthase that also catalyzes the synthesis of 3OHC_8_-HSL (12, 13, 16), as well as the BtaR3 transcriptional regulator, the product of the *btaR*3 gene located next to *btaI*3.

The main goal of this study was to dissect the QS regulatory network of *B. thailandensis* E264 to reveal the interactions existing between the QS-1, QS-2, and QS-3 systems. Besides verifying previously proposed and established interactions, we uncovered several interconnections between the QS-1, QS-2, and QS-3 systems, providing a comprehensive picture of the complex QS network in *B. thailandensis* E264. Ultimately, this study will contribute to a better appreciation of the QS regulatory mechanism of gene expression in *B. thailandensis*, and in particular those related to pathogenicity in *B. pseudomallei*.

## Materials and methods

### Bacterial strains and culture conditions

The bacterial strains used in this study are listed in **Table S1**. Unless otherwise stated, all bacteria were cultured at 37°C in Tryptic Soy Broth (TSB; BD Difco™, Mississauga, ON, Canada), with shaking (240 rpm) in a TC-7 roller drum (New Brunswick, Canada), or on Petri dishes containing TSB solidified with 1.5% agar. When required, antibiotics were used at the following concentrations: 15 μg/mL tetracycline (Tc) and 25 μg/mL gentamycin (Gm) for *Escherichia coli* DH5α, while Tc was used at 200 μg/mL for *B. thailandensis* E264. All measurements of optical density (OD_600_) were acquired with a Thermo Fisher Scientific NanoDrop® ND-1000 Spectrophotometer.

### Construction of plasmids

All plasmids used in this study are described in **Table S2**. Amplification of the promoter regions of *btaI*1, *btaI*2, and *btaI*3 was performed from genomic DNA of *B. thailandensis* E264 using appropriate primers (**Table S3**). The amplified products were digested with the FastDigest restriction enzymes *Xho*I and *Bam*HI (Thermo Fisher Scientific) and inserted by T4 DNA ligase (Bio Basic, Inc., Markham, ON, Canada) within the corresponding restriction sites in the mini-CTX-*lux* plasmid (20), generating the transcriptional reporters pSLG02, pSLG03, and pSLG04, respectively. Amplification of *btaR*1 and *btaR*3 was similarly performed using the primers shown in **Table S3** and the products were digested with the restriction enzymes *Xba*I and *Sac*I before ligation within the corresponding restriction sites in the pJN105 plasmid (21), generating the arabinose-inducible expression vectors pSLG23 and pSLG24, respectively. All primers were from Alpha DNA (Montreal, QC, Canada).

### Reporter strains construction

The mini-CTX-*btaI*1-*lux*, mini-CTX-*btaI*2-*lux*, and mini-CTX-*btaI*3-*lux* transcriptional reporters were integrated into the chromosome of *B. thailandensis* E264 strains through conjugation with *E. coli* χ7213 followed by selection with Tc. Successful chromosomal insertion of the *btaI*1-*lux*, *btaI*2-*lux*, and *btaI*3-*lux* plasmids was confirmed by PCR using appropriate primers.

### LC-MS/MS quantification of AHLs

The concentration of AHLs was determined from culture samples of *B. thailandensis* E264 obtained at different time points during bacterial growth, by liquid chromatography coupled to tandem mass spectrometry (LC-MS/MS). The samples were prepared and analyzed as described previously (22). 5,6,7,8-tetradeutero-4-hydroxy-2-heptylquinoline (HHQ-d4) was used as an internal standard. All experiments were performed in triplicate and carried out at least twice independently.

### Measurement of *btaI*1-*lux*, *btaI*2-*lux*, and *btaI*3-*lux* reporters’ activity

Expression from the promoter regions of *btaI*1, *btaI*2, or *btaI*3 was quantified by measuring the luminescence of *B. thailandensis* E264 cultures carrying the corresponding chromosomal reporters. Overnight bacterial cultures were diluted in TSB to an initial OD_600_ = 0.1 and incubated as indicated above. The luminescence was regularly determined from culture samples using a multi-mode microplate reader (Cytation^TM^ 3, BioTek Instruments, Inc., Winooski, VT, USA) and expressed in relative light units per culture optical density (RLU/OD_600_). For experiments with additions of AHLs, cultures were supplemented or not with 10 μM C_8_-HSL, 3OHC_10_-HSL, and 3OHC_8_-HSL (Sigma-Aldrich Co., Oakville, ON, Canada) from stocks prepared in HPLC-grade acetonitrile. Acetonitrile only was added in controls. All experiments were performed with three biological replicates and repeated at least twice.

### Heterologous *E. coli* expression systems for BtaR1, BtaR2, and BtaR3 regulation of *btaI*1, *btaI*2, and *btaI*3 expressions

Recombinant *E. coli* DH5α containing both a transcriptional fusion *btaI*1-*lux*, *btaI*2-*lux*, or *btaI*3-*lux* and an arabinose-inducible expression vector pJN105-*btaR*1, pJN105-*btaR*2, or pJN105-*btaR*3 were used to determine the response of the *btaI*1, *btaI*2, and *btaI*3 promoters genes to the BtaR1, BtaR2, and BtaR3 transcriptional regulators. Overnight bacterial cultures of *E. coli* DH5α carrying the respective plasmid combinations were diluted in LB broth (Alpha Biosciences, Inc., Baltimore, MD, USA), with appropriate antibiotics and grown in triplicate at 37°C, with shaking in a TC-7 roller drum. When reaching an OD_600_ = 0.5, cultures were supplemented with 1 or 10 μM C_8_-HSL, 3OHC_10_-HSL, or 3OHC_8_-HSL. Acetonitrile only was added in controls. The BtaR1, BtaR2, and BtaR3 expression vectors were induced with 0.2% L-arabinose (w/v). The *btaI*1-*lux*, *btaI*2-*lux,* and *btaI*3-*lux* luciferase activity was measured every 30 min during 10 hrs as described above. All experiments were repeated at least three times.

### Quantitative reverse-transcription PCR experiments

Total RNA of *B. thailandensis* E264 cultures at an OD_600_ = 4.0 was extracted with the PureZOL RNA Isolation Reagent (Bio-Rad Laboratories, Mississauga, ON, Canada) and treated twice with the TURBO DNA-free^TM^ Kit (Ambion Life Technologies, Inc., Burlington, ON, Canada), according to the manufacturer’s instructions. Extractions were done on three different bacterial cultures. Quality and purity controls were confirmed by agarose gel electrophoresis and UV spectrophotometric analysis, respectively. cDNA synthesis was performed using the iScript^TM^ Reverse Transcription Supermix (Bio-Rad Laboratories) and amplification was accomplished on a Corbett Life Science Rotor-Gene® 6000 Thermal Cycler, using the SsoAdvanced^TM^ Universal SYBR® Green Supermix (Bio-Rad Laboratories), according to the manufacturer’s protocol. The reference gene was *ndh* (23). All primers used for cDNA amplification are presented in **Table S4**. Gene expression differences between *Burkholderia thailandensis* E264 strains were calculated using the 2^(-ΔΔ(CT))^ formula (24). A threshold of 0.5 was chosen as significant. All experiments were performed in triplicate and carried out at least twice independently.

## Results

### The *B. thailandensis* QS-1, QS-2, and QS-3 systems are sequentially activated

*B. thailandensis* E264 produces 3OHC_10_-HSL and to lesser extents, C_8_-HSL and 3OHC_8_-HSL (12, 15). Considering that non-simultaneous production of AHLs was suggested in *B. pseudomallei* (16), we hypothesized that these three AHLs are differentially produced over the bacterial growth stages of *B. thailandensis* E264. We thus determined the production profiles of C_8_-HSL, 3OHC_10_-HSL, and 3OHC_8_-HSL at various time points of the bacterial growth. LC-MS/MS was used to quantify concentrations of these AHLs in *B. thailandensis* E264 wild-type cultures. We found that 3OHC_10_-HSL levels increased rapidly through the early logarithmic (OD_600_ ≈ 3.0) and late exponential phases (OD_600_ ≈ 5.0), but decreased later on (**Fig. 1A**). Interestingly, 3OHC_8_-HSL concentrations kept increasing all along bacterial growth to levels similar to 3OHC_10_-HSL (**Fig. 1A**), while C_8_-HSL only accumulated during the early exponential growth phase (OD_600_ ≈ 4.0), but then remained stable during the stationary phase (OD_600_ ≈ 8.0; **Fig. 1A**).

**Figure 1.**
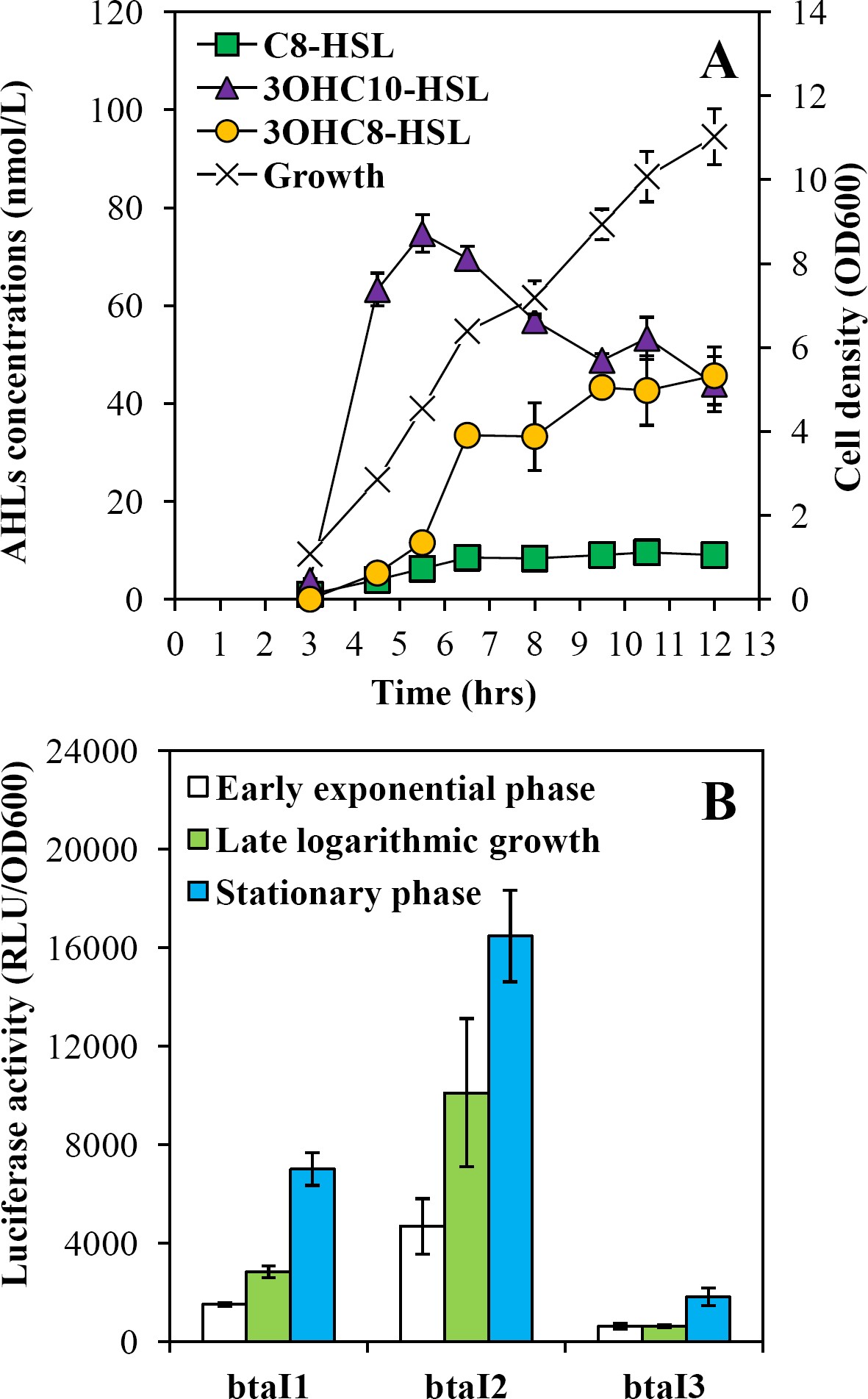
The QS-1, QS-2, and QS-3 systems are consecutively activated. (A) AHLs production was measured by LC-MS/MS during the different stages of bacterial growth in cultures of the wild-type strain of *B. thailandensis* E264. The error bars represent the standard deviation of the average for three replicates. (B) The luciferase activity of the *btaI*1-*lux*, *btaI*2-*lux*, and *btaI*3-*lux* chromosomal reporters was monitored during the early exponential phase (OD_600_ ≈ 3.0), the late logarithmic growth (OD_600_ ≈ 5.0), and the stationary phase (OD_600_ ≈ 8.0). The luminescence is expressed in relative light units per culture optical density (RLU/OD_600_).

To gain additional insights, AHLs biosynthesis was correlated to *btaI*1, *btaI*2, and *btaI*3 expressions. The activity of the chromosomal *btaI*1-*lux*, *btaI*2-*lux*, and *btaI*3-*lux* transcriptional reporters was measured during bacterial growth. In agreement with the AHLs production profiles, activation of both *btaI*1 and *btaI*2 was observed from the beginning of the exponential phase, whereas *btaI*3 was not activated until the stationary phase was reached (**Fig. 1B**). Collectively, our results point toward a sequential activation of the different QS systems in *B. thailandensis* E264.

### Interconnections between the three QS systems are observed

In order to verify whether the sequential activation of the QS-1, QS-2, and QS-3 systems results from interactions between these systems, we determined the AHLs production kinetics in cultures of the Δ*btaR*1 (JBT107), Δ*btaR*2 (JBT108), and Δ*btaR*3 (JBT109) mutants *versus* wild-type *B. thailandensis* E264. We also measured expressions of the AHL synthase-coding genes *btaI*1, *btaI*2, and *btaI*3 in the same background harboring a chromosomal *btaI*1-*lux*, *btaI*2-*lux*, or *btaI*3-*lux* transcriptional fusion.

BtaI1 produces C_8_-HSL and BtaR1 is considered the main regulator of *btaI*1 expression (12). Therefore, we were surprised to see an increased production of C_8_-HSL in the Δ*btaR*1 mutant in comparison with the wild-type strain (**Fig. 2A**). This overproduction in the absence of BtaR1 was principally detected from the end of the exponential phase. Nevertheless, transcription of the *btaI*1 gene was indeed lower in Δ*btaR*1 throughout the different stages of bacterial growth and was almost not detected in the early logarithmic growth (**Fig. 2B**). Following those results, it was important to confirm that *btaI*1 expression is directly modulated by BtaR1 in conjunction with C_8_-HSL. We monitored *btaI*1 expression in response to exogenous addition of C_8_-HSL in the wild-type strain of *B. thailandensis* E264 and in its Δ*btaR*1, Δ*btaI*1 (JBT101), and Δ*btaI*1Δ*btaI*2Δ*btaI*3 (JBT112) mutants. The *btaI*1 gene exhibited a comparable transcriptional profile in the absence of BtaR1 or C_8_-HSL, supporting that BtaR1/C_8_-HSL indeed activates *btaI*1 transcription (**Fig. S1A**). Accordingly, adding exogenous C_8_-HSL to the culture of the Δ*btaI*1 mutant as well as in the Δ*btaI*1Δ*btaI*2Δ*btaI*3 mutant restored *btaI*1 transcription (**Fig. S1A**). While *btaI*1 expression was induced in the wild-type strain culture supplemented with exogenous C_8_-HSL, no difference was noticed for the Δ*btaR*1 mutant (**Fig. S1A**). To confirm that *btaI*1 is directly activated by BtaR1, we measured the luciferase activity of *btaI*1*-lux* in the heterologous system *E. coli* DH5α, expressing *btaR*1 controlled by an arabinose-inducible promoter. In agreement with the detection of a putative *lux*-box sequence found in the promoter region of *btaI*1, BtaR1 increased *btaI*1 transcription (**Fig. S1B**). However, no significant further activation was observed in response to addition of C_8_-HSL (**Fig. S1B**). Interestingly, C_8_-HSL concentrations were also increased in the Δ*btaR*2 mutant, with a matching upregulation of *btaI*1 expression (**Fig. 2**). While C_8_-HSL was also overproduced in the absence of BtaR3 during the stationary phase, *btaI*1 transcription was downregulated in the Δ*btaR*3 mutant in comparison with the wild-type strain (**Fig. 2**). Collectively, these results suggest that *btaI*1 expression is also controlled by BtaR2 and BtaR3. Nevertheless, no direct interactions between the *btaI*1 promoter and neither BtaR2 nor BtaR3 were observed in our heterologous expression systems (data not shown).

**Figure 2.**
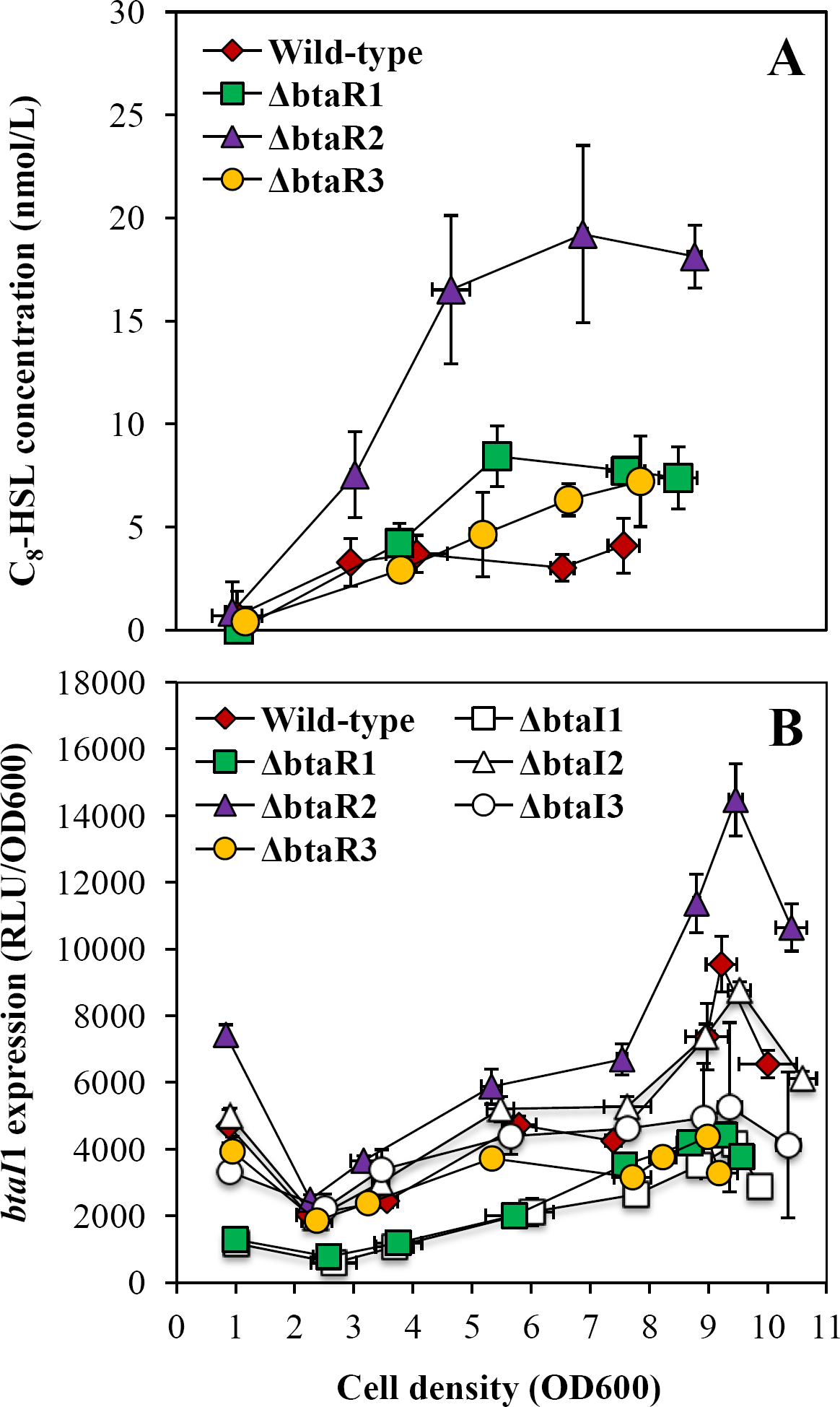
C_8_-HSL production and expression from the *btaI*1 promoter in the wild-type and the QS mutant strains of *B. thailandensis* E264. (A) Biosynthesis of C_8_-HSL was quantified using LC-MS/MS at various times during growth in cultures of the wild-type and of the Δ*btaR*1, Δ*btaR*2, and Δ*btaR*3 mutant strains of *B. thailandensis* E264. The error bars represent the standard deviation of the average for three replicates. The luminescence of the mini-CTX-*btaI*1-*lux* transcriptional fusion was monitored in cultures of the wild-type and of the Δ*btaR*1, Δ*btaR*2, Δ*btaR*3, Δ*btaI*1, Δ*btaI*2, and Δ*btaI*3 mutant strains of *B. thailandensis* E264 carrying a chromosomal *btaI*1-*lux* transcriptional reporter. The luminescence is expressed in relative light units per culture optical density (RLU/OD_600_).

3OHC_10_-HSL is produced by the BtaI2 synthase (12, 15). While BtaR2 directly activates *btaI*2 expression in response to 3OHC_10_-HSL and 3OHC_8_-HSL, the latter also produced by BtaI2 (12, 15), the impact of BtaR2 on the production of these two AHLs is still unknown. We observed that both 3OHC_10_-HSL biosynthesis and *btaI*2 expression were almost completely abolished in the Δ*btaR*2 mutant (**Fig. 3**), confirming that BtaR2 is the main regulator of 3OHC_10_-HSL biosynthesis via its effect on *btaI*2 expression. Despite the absence of BtaR2, we detected a slight, but consistent and highly reproducible, production of 3OHC_10_-HSL during the stationary phase (**Fig. 3A**). Accordingly, transcription of *btaI*2 was also slightly augmented later (**Fig. 3B**). Interestingly, while 3OHC_10_-HSL concentrations were strongly increased in the Δ*btaR*1 mutant in comparison with the wild-type strain (**Fig. 3A**), expression of *btaI*2 was not higher in the absence of BtaR1 (**Fig. 3B**). 3OHC_10_-HSL concentrations were also increased in the Δ*btaR*3 mutant background (**Fig. 3A**). The effect of BtaR3’s absence on 3OHC_10_-HSL production was only observed from the end of logarithmic growth (**Fig. 3A**), and loss of BtaR3 did not affect *btaI*2 transcription either (**Fig. 3B**). Collectively, these observations indicate that, in addition to BtaR2, both BtaR1 and BtaR3 also influence the biosynthesis of 3OHC_10_-HSL. Nevertheless, no discernible difference in *btaI*2 transcription was observed in the absence of BtaR1 and BtaR3, suggesting that the effect of these transcriptional regulators on the QS-2 system is indirect.

**Figure 3.**
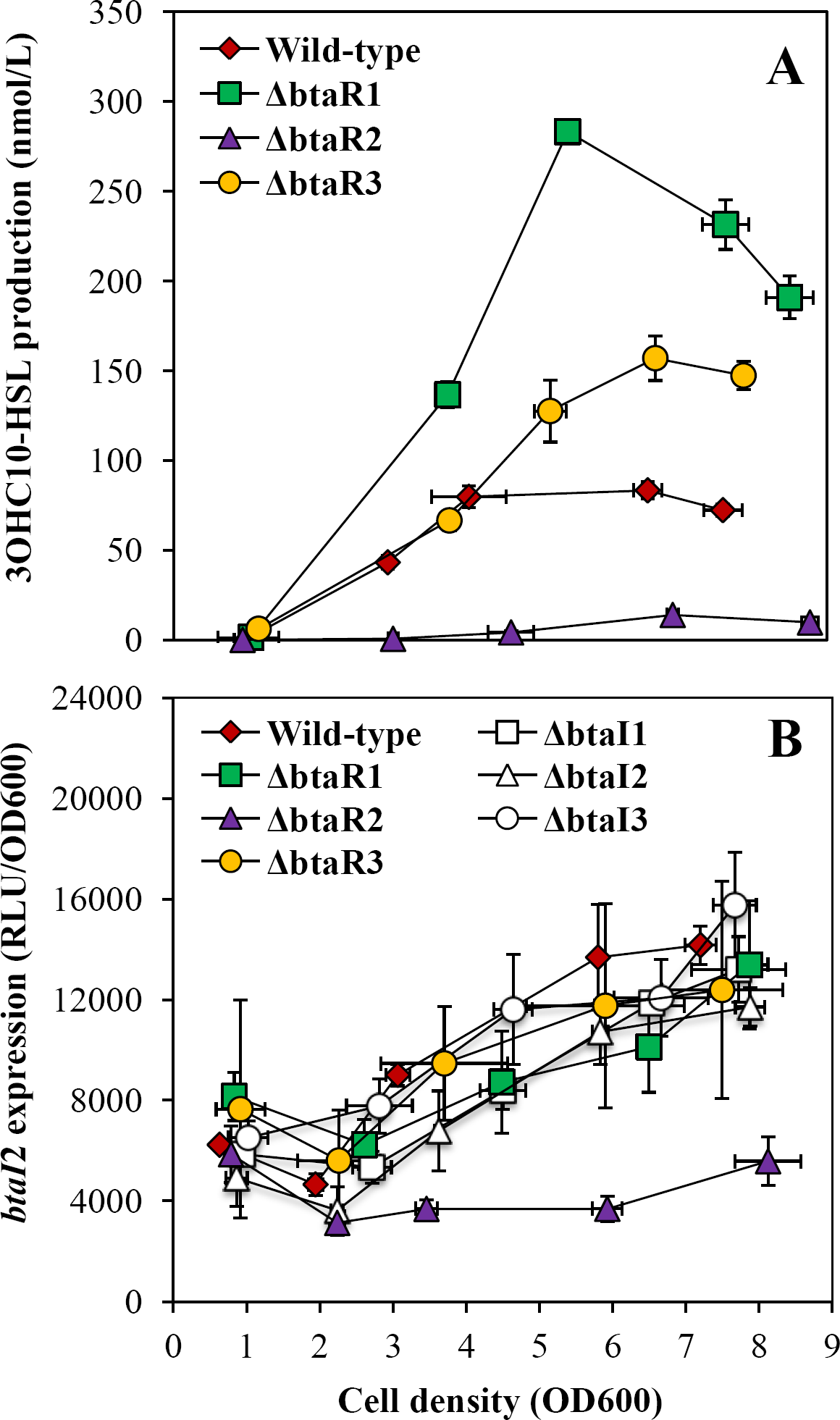
3OHC_10_-HSL production and expression from the *btaI*2 promoter in the wild-type and the QS mutant strains of *B. thailandensis* E264. (A) Biosynthesis of 3OHC_10_-HSL was quantified using LC-MS/MS at various times during growth in cultures of the wild-type and of the Δ*btaR*1, Δ*btaR*2, and Δ*btaR*3 mutant strains of *B. thailandensis* E264. The error bars represent the standard deviation of the average for three replicates. The luminescence of the mini-CTX-*btaI*2-*lux* transcriptional fusion was monitored in cultures of the wild-type and of the Δ*btaR*1, Δ*btaR*2, Δ*btaR*3, Δ*btaI*1, Δ*btaI*2, and Δ*btaI*3 mutant strains of *B. thailandensis* E264 carrying a chromosomal *btaI*2-*lux* transcriptional reporter. The luminescence is expressed in relative light units per culture optical density (RLU/OD_600_).

BtaI3 is mainly responsible for 3OHC_8_-HSL biosynthesis (12, 15). While no discernible difference in 3OHC_8_-HSL concentrations was detected in cultures of the Δ*btaR*3 mutant when compared to the wild-type strain, the levels of *btaI*3 transcription were decreased in the absence of BtaR3 (**Fig. 4B**). To confirm whether transcription of *btaI*3 is dependent on BtaR3, as well as on 3OHC_8_-HSL, *btaI*3 expression was measured in the wild-type strain and in the Δ*btaR*3 and Δ*btaI*3 (JBT103) mutants supplemented or not with exogenous 3OHC_8_-HSL. We found that *btaI*3 was similarly downregulated in these two backgrounds, suggesting that BtaR3 activates *btaI*3 in response to 3OHC_8_-HSL (**Fig. S2A**). Accordingly, *btaI*3 transcription was not affected by the addition of 3OHC_8_-HSL in the Δ*btaR*3 mutant, but was increased in the wild-type strain culture under the same conditions (**Fig. S2A**). Unexpectedly, adding exogenous 3OHC_8_-HSL to the culture of the Δ*btaI*3 mutant did not restore *btaI*3 transcription to wild-type levels (**Fig. S2A**). Since 3OHC_8_-HSL production is not completely abolished in the Δ*btaI*3 mutant strain (12, 13, 16), we examined the effect of 3OHC_8_-HSL addition on *btaI*3 expression in the AHL-defective Δ*btaI*1Δ*btaI*2Δ*btaI*3 mutant and indeed observed that transcription of *btaI*3 was restored by 3OHC_8_-HSL in this background (**Fig. S2A**). To gain insight into the regulation of *btaI*3, the activity of *btaI*3*-lux* was also quantified in *E. coli* DH5α, containing a BtaR3 expression vector with an arabinose-inducible promoter. While no putative *lux*-box sequence was found in the promoter region of *btaI*3, we observed an increase in *btaI*3 transcription in the presence of BtaR3 (**Fig. S2B**). However, we did not see any effect of 3OHC_8_-HSL addition (**Fig. S2B**). Taken together, these data suggest that *btaI*3 expression is controlled by additional AHLs and/or alternative LuxR-type transcriptional regulators.

**Figure 4.**
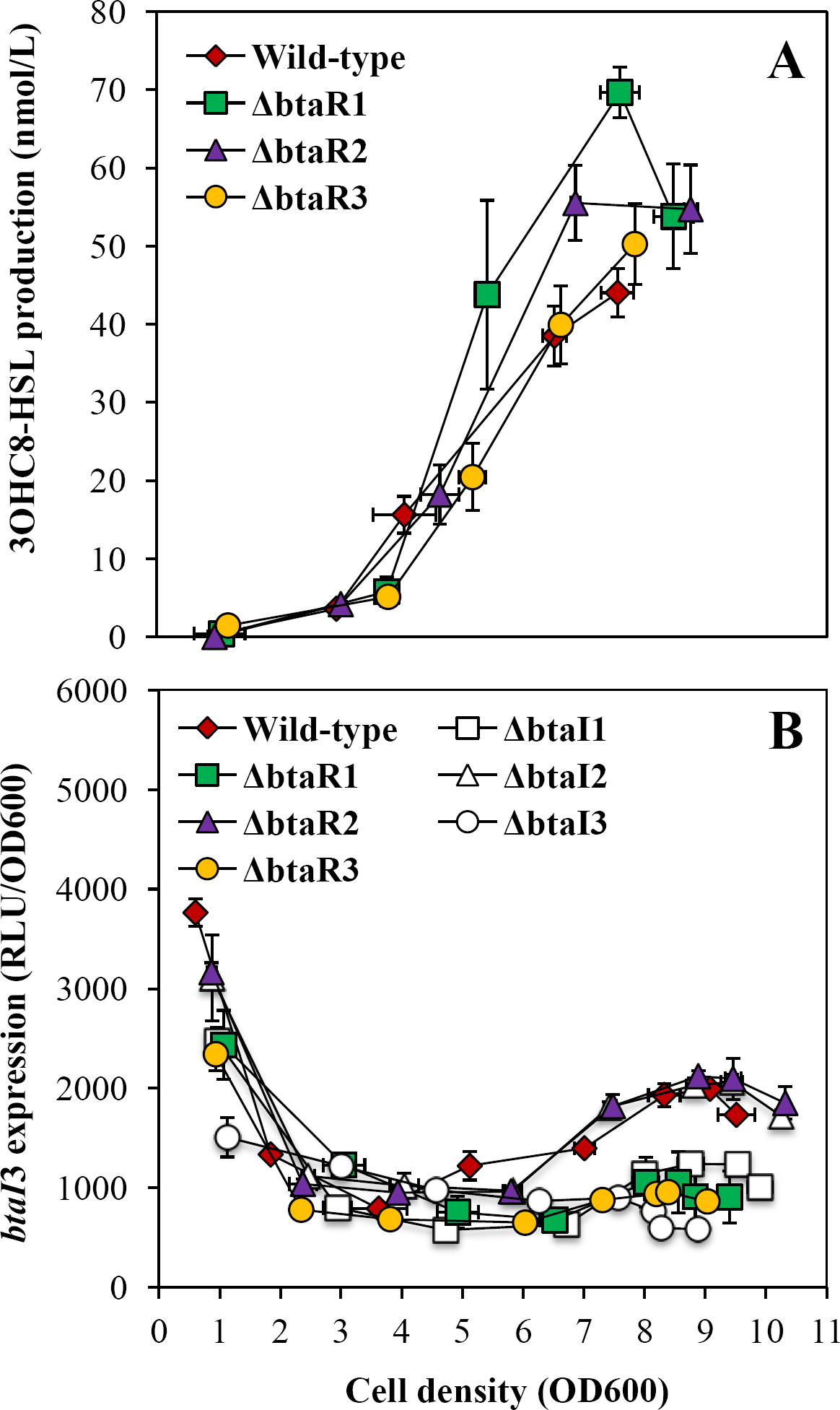
3OHC_8_-HSL production and expression from the *btaI*3 promoter in the wild-type and the QS mutant strains of *B. thailandensis* E264. (A) Biosynthesis of 3OHC_8_-HSL was quantified using LC-MS/MS at various times during growth in cultures of the wild-type and of the Δ*btaR*1, Δ*btaR*2, and Δ*btaR*3 mutant strains of *B. thailandensis* E264. The error bars represent the standard deviation of the average for three replicates. The luminescence of the mini-CTX-*btaI*3-*lux* transcriptional fusion was monitored in cultures of the wild-type and of the Δ*btaR*1, Δ*btaR*2, Δ*btaR*3, Δ*btaI*1, Δ*btaI*2, and Δ*btaI*3 mutant strains of *B. thailandensis* E264 carrying a chromosomal *btaI*3-*lux* transcriptional reporter. The luminescence is expressed in relative light units per culture optical density (RLU/OD_600_).

As previously noted for C_8_-HSL and 3OHC_10_-HSL, the levels of 3OHC_8_-HSL were enhanced in the Δ*btaR*1 mutant in comparison with the wild-type strain (**Fig. 4A**). While 3OHC_10_-HSL overproduction was observed during the different stages of bacterial growth (**Fig. 3A**), augmentation of 3OHC_8_-HSL concentrations, similarly to C_8_-HSL, principally occurred from the end of the exponential phase in the Δ*btaR*1 mutant (**Fig. 4A**). As seen for *btaI*1, expression of *btaI*3 was surprisingly lower in the absence of BtaR1 (**Fig. 4B**). Additionally, we observed an increase in the amounts of 3OHC_8_-HSL in the Δ*btaR*2 mutant from the stationary phase (**Fig. 4A**). Nevertheless, no obvious change in expression of *btaI*3 was visible (**Fig. 4B**), suggesting that unknown factors are involved in the regulation of 3OHC_8_-HSL biosynthesis.

We also analysed AHLs production in the Δ*btaR*4 (JBT110) and Δ*btaR*5 (JBT111) mutants and no difference with the wild-type strain production was found (data not shown).

### BtaR1, BtaR2, and BtaR3 are transcriptionally intertwined

In order to verify whether the QS modulatory cascade also involves cross-regulations between BtaR1, BtaR2, and BtaR3, *btaR*1, *btaR*2, and *btaR*3 expressions were assessed by quantitative reverse-transcription PCR (qRT-PCR) in the *B. thailandensis* E264 wild-type strain and in the AHL-defective Δ*btaI*1Δ*btaI*2Δ*btaI*3 mutant during the exponential phase (OD_600_ ≈ 4.0). Interestingly, expressions of all transcriptional regulators were affected by the absence of AHLs (**Fig. 5**), indicating that *btaR*1, *btaR*2, and *btaR*3 are QS-controlled. While *btaR*1 transcription was increased in the Δ*btaI*1Δ*btaI*2Δ*btaI*3 mutant when compared to the wild-type strain (**Fig. 5A**), *btaR*2 and *btaR*3 were both downregulated in the absence of AHLs (**Figs. 5B and C**). To further investigate the impact of AHLs on *btaR*1, *btaR*2, and *btaR*3 expressions, their transcription was measured in the Δ*btaI*1Δ*btaI*2Δ*btaI*3 mutant supplemented with exogenous C_8_-HSL, 3OHC_10_-HSL, or 3OHC_8_-HSL. Interestingly, *btaR*1, *btaR*2, and *btaR*3 expressions were restored in the presence of AHLs (**Fig. 5**), suggesting that these genes are transcriptionally intertwined. Collectively, our results indicate that the QS-1, QS-2, and QS-3 systems interdependence also implicates cross-modulations between BtaR1, BtaR2, and BtaR3.

**Figure 5.**
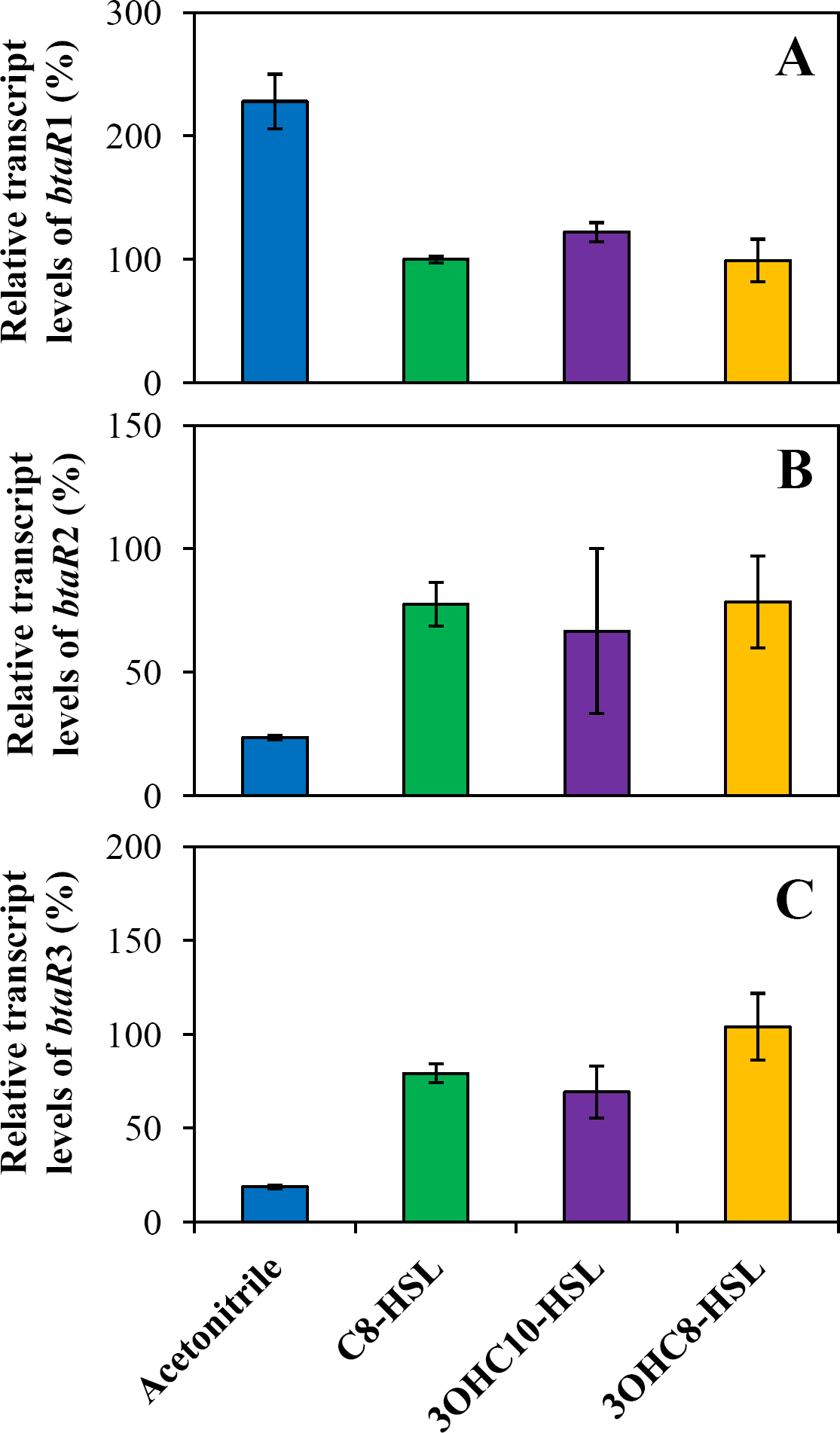
Effects of AHLs on the expressions of *btaR*1, *btaR*2, and *btaR*3. The relative transcript levels of (A) *btaR*1, (B) *btaR*2, and (C) *btaR*3 from the *B. thailandensis* E264 wild-type and its Δ*btaI*1Δ*btaI*2Δ*btaI*3 mutant strain was estimated by qRT-PCR experiments. Cultures were supplemented with 10 μM C_8_-HSL, 3OHC_10_-HSL, or 3OHC_8_-HSL. Acetonitrile only was added in controls. The results are presented as relative quantification of genes transcription compared to the wild-type normalized to 100%. The error bars represent the standard deviation of the average for three replicates.

### Expressions of *btaI*1, *btaI*2 and *btaI*3 are modulated by AHLs

To further elucidate the regulatory mechanisms directing *btaI*1, *btaI*2, and *btaI*3 expressions, the activity of the chromosomal *btaI*1-*lux*, *btaI*2-*lux*, and *btaI*3-*lux* transcriptional reporters was measured in the AHL-defective Δ*btaI*1Δ*btaI*2Δ*btaI*3 mutant supplemented or not with exogenous AHLs. We had noted that the QS-1 and QS-2 systems were both activated from the logarithmic growth, whereas activation of the QS-3 system started in the stationary phase (**Fig. 1**). Therefore, experiments with *btaI*1-*lux* and *btaI*2-*lux* were done during the exponential phase (OD_600_ ≈ 4.0), while those with *btaI*3-*lux* were performed during the stationary phase (OD_600_ ≈ 8.0). Furthermore, AHLs impact on *btaI*1, *btaI*2, and *btaI*3 expressions was also estimated by monitoring the activity of *btaI*1-*lux*, *btaI*2-*lux*, and *btaI*3-*lux* in cultures of the Δ*btaI*1, Δ*btaI*2 (JBT102), and Δ*btaI*3 mutants *versus* wild-type *B. thailandensis* E264.

We have demonstrated that transcription of *btaI*1 is directly controlled by BtaR1 and activated by C_8_-HSL (**Fig. S1**). Additionally, *btaI*1 expression was enhanced in the presence of 3OHC_10_-HSL or 3OHC_8_-HSL in the AHL-negative mutant background (**Fig. 6A**). Using a heterologous expression system, we also confirmed that BtaR2 modulates directly *btaI*2 transcription in response to 3OHC_10_-HSL or 3OHC_8_-HSL (**Fig. S4**). While *btaI*2 expression was accordingly activated by both 3OHC_10_-HSL and 3OHC_8_-HSL in the Δ*btaI*1Δ*btaI*2Δ*btaI*3 mutant (**Fig. 6B**), we noticed again that transcription of *btaI*2 was most strongly increased by the presence of 3OHC_10_-HSL (**Fig. 6B**). Transcription of *btaI*3 was at least doubled in the Δ*btaI*1Δ*btaI*2Δ*btaI*3 mutant strain culture when supplemented with any of the three AHLs (**Fig. 6C**). Here, *btaI*3 expression was substantially increased in the presence of 3OHC_8_-HSL (**Fig. 6C**). Since all three AHLs seem able to activate *btaI*3 expression, we investigated whether their respective influence changes over the various growth phases. Strikingly, activation of *btaI*3 transcription by C_8_-HSL is more prominent during the logarithmic growth phase, whereas *btaI*3 is mostly activated by 3OHC_8_-HSL during the stationary phase (**Fig. 7**).

**Figure 6.**
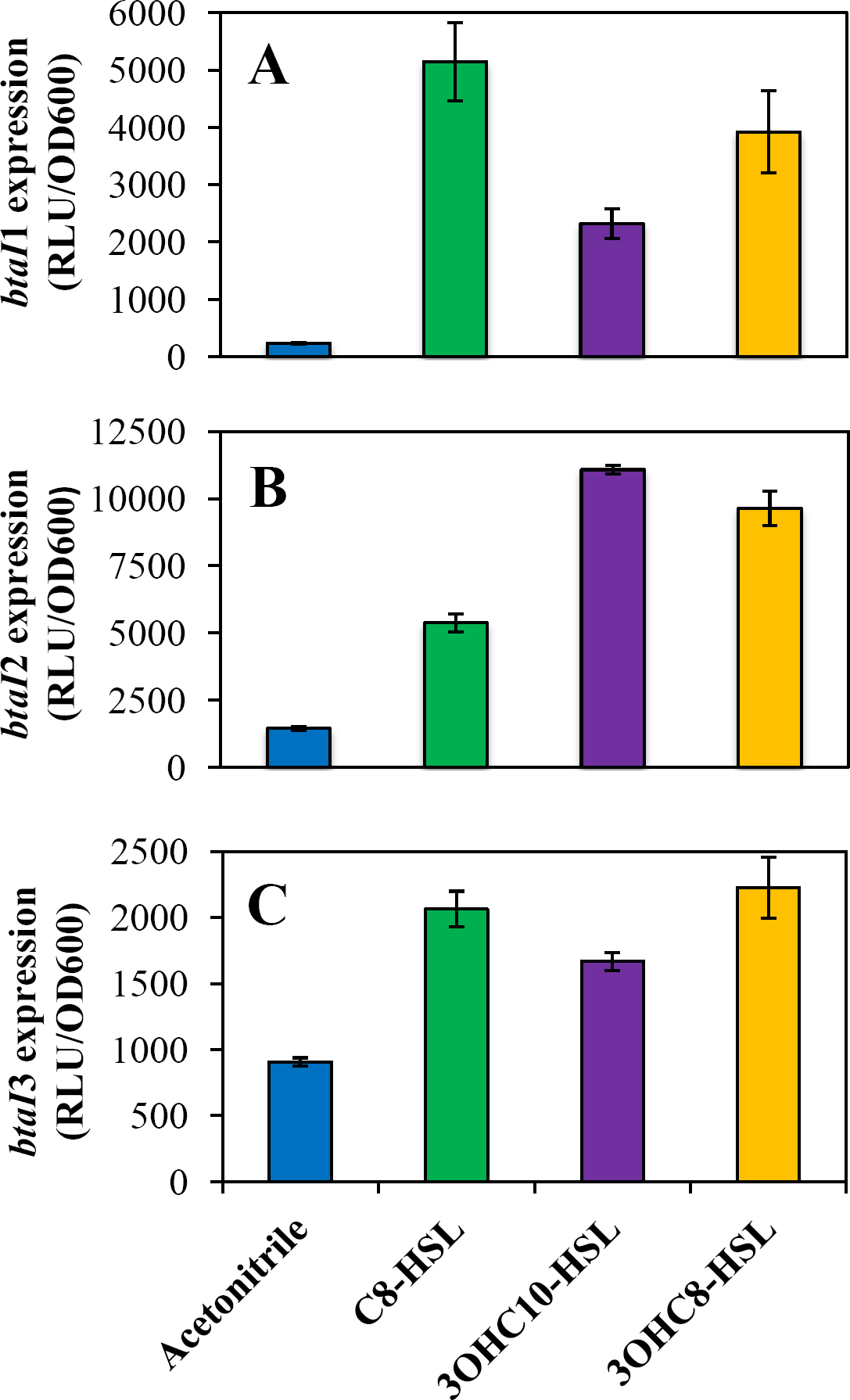
Activation of expression from the *btaI*1, *btaI*2, and *btaI*3 promoters by AHLs. The luminescence of the mini-CTX-*btaI*1-*lux*, mini-CTX-*btaI*2-*lux*, and mini-CTX-*btaI*3-*lux* transcriptional fusions was monitored in cultures of the *B. thailandensis* E264 Δ*btaI*1Δ*btaI*2Δ*btaI*3 mutant strain haboring a chromosomal (A) *btaI*1-*lux*, (B) *btaI*2-*lux*, and (C) *btaI*3-*lux* transcriptional reporter, respectively. Cultures were supplemented with 10 μM C_8_-HSL, 3OHC_10_-HSL, or 3OHC_8_-HSL. Acetonitrile only was added in controls. The error bars represent the standard deviation of the average for three replicates. The luminescence is expressed in relative light units per culture optical density (RLU/OD_600_).

**Figure 7.**
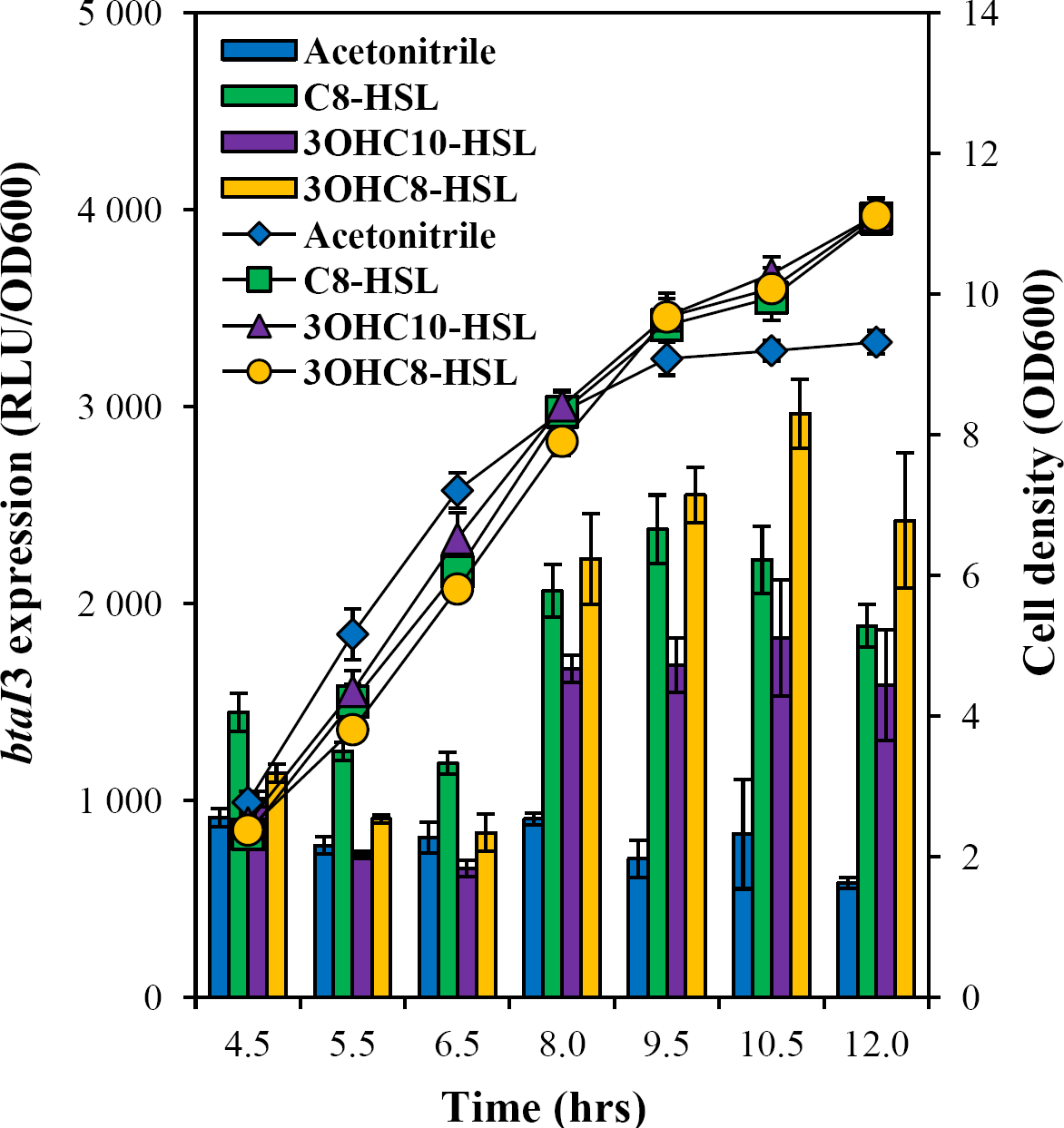
Activation of expression from the *btaI*3 promoter by AHLs. The luciferase activity (bars) of the mini-CTX-*btaI*3-*lux* transcriptional fusion was monitored at various times during growth (lines) in cultures of the *B. thailandensis* E264 Δ*btaI*1Δ*btaI*2Δ*btaI*3 mutant, haboring a chromosomal *btaI*3-*lux* transcriptional reporter. Cultures were supplemented with 10 μM C_8_-HSL, 3OHC_10_-HSL, or 3OHC_8_-HSL. Acetonitrile only was added in controls. The error bars represent the standard deviation of the average for three replicates. The luminescence is expressed in relative light units per culture optical density (RLU/OD_600_).

## Discussion

Although the QS-1, QS-2, and QS-3 systems of *B. thailandensis* had been previously described (12, 15), a detailed picture of the interactions composing this complex QS regulatory network had never been exposed.

As previously described for *B. pseudomallei* KHW (16), we observed variations in the biosynthesis of the main AHLs produced by *B. thailandensis* E264 throughout the bacterial growth phases (**Fig. 1A**), as well as in the transcription of the three AHL synthase-coding genes *btaI*1, *btaI*2, and *btaI*3 (**Fig. 1B**). These observations highlighted the timing of expression of QS-1, QS-2, and QS-3 during the different stages of growth and consequently the existence of potential interactions between these QS-systems. While C_8_-HSL is generally considered the primary AHL produced by *Burkholderia* spp. (4), and is indeed predominately detected in stationary phase cultures of *B. pseudomallei* K96243 and *B. mallei* ATCC 23344 (14, 16), we confirmed that 3OHC_10_-HSL is actually the most abundant AHL found in *B. thailandensis* E264 cultures during the different stages of growth (**Fig. 1A**), revealing the importance of the QS-2 system in the QS circuitry of *B. thailandensis* E264 (**Fig. 8**). While 3OHC_10_-HSL and 3OHC_8_-HSL are both reported to be produced by BtaI2 (12, 15), we observed differences in the production profiles of these AHLs (**Fig. 1A**). Indeed, the concentration of 3OHC_10_-HSL first increased and then decreased in the exponential and stationary phases respectively, whereas the levels of 3OHC_8_-HSL increased throughout the different stages of growth similarly to the expression pattern of *btaI*2 (**Fig. 1B**). This suggests that 3OHC_8_-HSL is produced by BtaI2 at the expense of 3OHC_10_-HSL. Since BtaI3 was also shown to catalyze the biosynthesis of 3OHC_8_-HSL (12, 15), and considering the *btaI*3 transcription profile, we assume that production of 3OHC_8_-HSL during the exponential phase is mainly the product of BtaI2, whereas BtaI3 is principally responsible for 3OHC_8_-HSL biosynthesis during the stationary phase, implying a cooperation between the QS-2 and QS-3 systems (**Fig. 8**). Consistently, *btaI*2 activation by 3OHC_8_-HSL was detected in the exponential phase (**Fig. 6B**), whereas we determined that 3OHC_8_-HSL activated *btaI*3 in the stationary phase (**Fig. 6C**). However, we observed that *btaI*2 expression was mainly enhanced by 3OHC_10_-HSL, indicating that BtaR2 exhibits a higher affinity for 3OHC_10_-HSL than for 3OHC_8_-HSL. The *bpsI*2 gene that codes for the BpsI2 synthase was also shown to be substantially activated by 3OHC_10_-HSL in *B. pseudomallei* KHW (16). Still, the levels of expression of *btaI*2 were similar in the wild-type strain of *B. thailandensis* E264 and in the Δ*btaI*2 mutant (**Fig. 3B**). Considering that 3OHC_8_-HSL is produced in the absence of BtaI2 (15), we must conclude that both 3OHC_10_-HSL and 3OHC_8_-HSL can induce the transcription of *btaI*2. Accordingly, Majerczyk *et al.* demonstrated that *btaR*2 expression is stimulated by 3OHC_8_-HSL (19). We determined that the transcription of *btaR*2 was affected by the absence of AHLs, indicating that *btaR*2 is also QS-controlled (**Fig. 5B**), and we observed an activation by both 3OHC_10_-HSL and 3OHC_8_-HSL, suggesting that *btaR*2 transcription is auto-regulated as well. Surprisingly, expressions of *btaI*2 and *btaR*2 were also induced by C_8_-HSL. Since we confirmed that BtaR2 does not specifically respond to C_8_-HSL (**Fig. S4**), we assume that activation by this AHL is linked to an alternative LuxR-type transcriptional regulator and that the transcription of *btaI*2 and *btaR*2 is not only under BtaR2 control.

**Figure 8.**
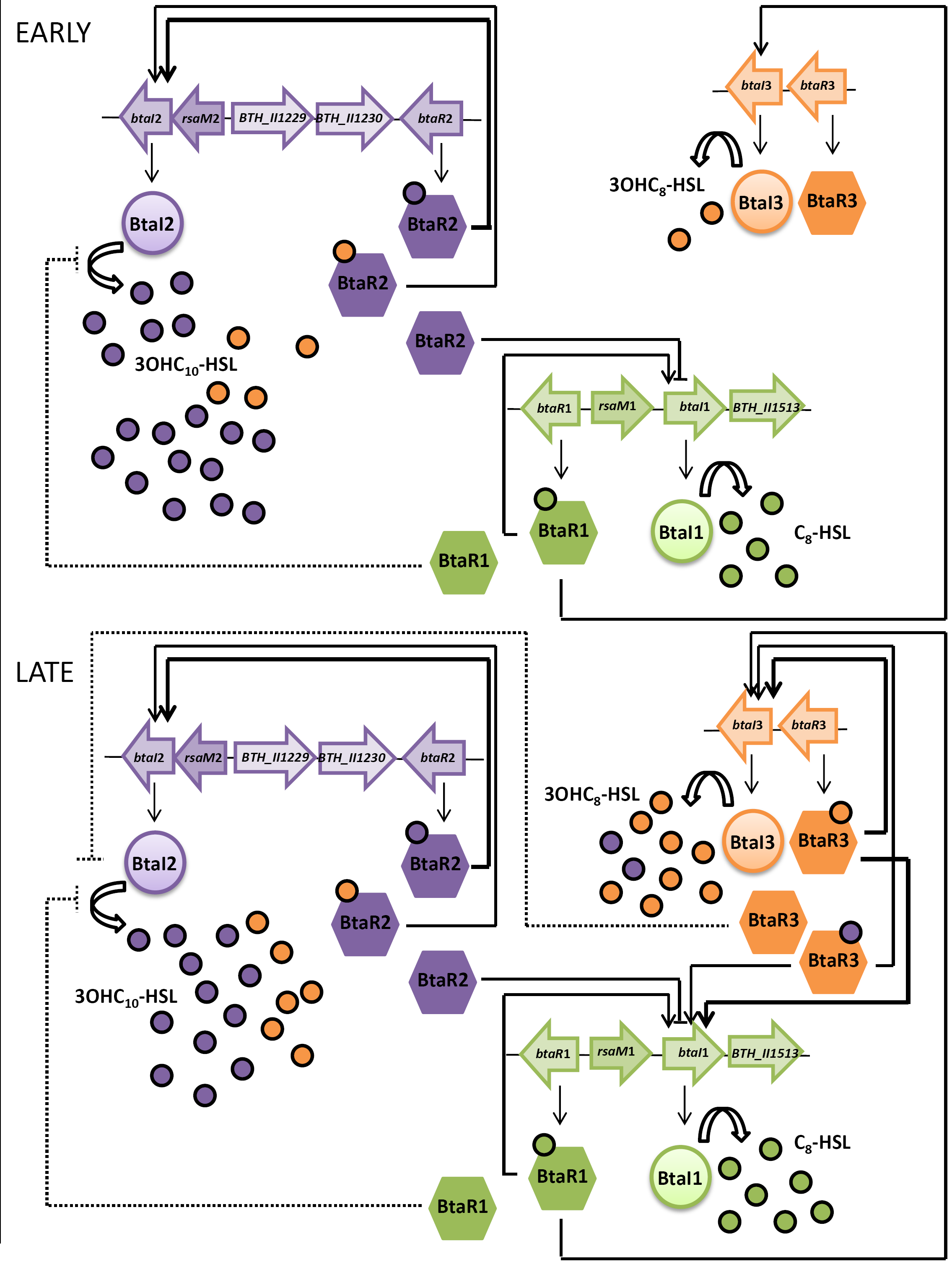
Proposed interactions between the QS-1, QS-2, and QS-3 systems.

Since the Δ*btaR*1 and Δ*btaR*3 mutants both accumulated 3OHC_10_-HSL principally in the exponential and stationary phases, respectively (**Fig. 3A**), suggesting that the regulation of the levels of 3OHC_10_-HSL does not only involve BtaR2 and might imply a dynamic coordination of the *B. thailandensis* E264 QS circuitry. The differences observed between the 3OHC_10_-HSL production and *btaI*2 transcription profiles could also be attributed to interactions between the QS-1, QS-2, and QS-3 systems. Nevertheless, neither BtaR1, nor BtaR3 affected the transcription of *btaI*2 during the different stages of growth (**Fig. 3B**). We thus hypothesize that BtaR1 and BtaR3 act indirectly on *btaI*2 expression by modulating the activity of BtaI2 for instance or that unknown factors might be involved in the regulation of 3OHC_10_-HSL biosynthesis. The *btaI*2 gene is predicted to be organized in operon with two additional genes, namely *BTH_II1226* (*btaE*) and *BTH_II1228* (*btaF*; **Fig. S3**). Interestingly, the *BTH_II1228* gene encodes a hypothetical protein conserved in the *Burkholderia* genus (25). This hypothetical protein is 37% identical to RsaM of *B. cenocepacia* J2315 (26). RsaM was shown to inhibit AHLs production in *Pseudomonas fuscovaginae* (27). While the *BTH_II1228* gene is located in a cluster involved in bactobolin biosynthesis (28), its involvement was actually not demonstrated. Interestingly, we observed that C_8_-HSL, 3OHC_10_-HSL, and 3OHC_8_-HSL concentrations are all increased when the *BTH_II1228* gene is inactivated (S. Le Guillouzer, M. C. Groleau, and E. Déziel, unpublished data). Therefore, we rename this hypothetical protein RsaM2.

Interestingly, a gene encoding a hypothetical protein sharing 63% identity with the *B. cenocepacia* J2315 RsaM that we call RsaM1 was also found between the *btaI*1 and *btaR*1 genes (**Fig. S3**). While a mutant in the *rsaM* gene of *B. cenocepacia* H111 showed higher levels of C_8_-HSL, expression of the QS regulatory *cepI* and *cepR* genes, encoding respectively the LuxI-type synthase CepI and the LuxR-type transcriptional regulator CepR, were lowered in the *rsaM*-mutant in comparison with the wild-type strain (26). In agreement with the putative *lux*-box sequence found in the promoter region of *rsaM*, the *rsaM* gene was shown to be positively and directly controlled by CepR (29). Investigating the effect of RsaM1 on AHLs biosynthesis in *B. thailandensis* E264 showed that C_8_-HSL and 3OHC_8_-HSL are both overproduced in the *rsaM*1-mutant when compared to the wild-type strain (S. Le Guillouzer, M. C. Groleau, and E. Déziel, unpublished data). How RsaM1 impacts *btaI*1 and *btaR*1 expressions has not been determined yet and is still under investigation. Interestingly, our transcriptomic sequencing analyses indicate that QS positively regulates the expression of the *rsaM*1 gene, as well as *rsaM*2 (S. Le Guillouzer, M. C. Groleau, and E. Déziel, unpublished data), and could thus be a target of BtaR1.

The QS-1 system, that consists in BtaI1 and BtaR1, is homologous to the *B. pseudomallei* BpsI/BpsR and *B. mallei* BmaI1/BpR1 QS systems. The BtaI1 protein of *B. thailandensis* E264 is 97% identical to *B. pseudomallei* K96243 BpsI and *B. mallei* ATCC 23344 BmaI1, whereas the BtaR1 protein shares 99% identity with BpsR and BmaR1. Chandler *et al*. (12) demonstrated that BtaI1 is responsible for C_8_-HSL production, as described previously for the BpsI and BmaI1 synthases (14, 30). Furthermore, the BpsR and BmaR1 transcriptional regulators were shown to directly activate the *bpsI* and *bmaI*1 genes in response to C_8_-HSL (14, 31). Accordingly, Majerczyk *et al*. reported that BtaR1 modulates positively *btaI*1 transcription (19). We observed a strong BtaR1-dependant induction of *btaI*1 by C_8_-HSL (**Fig. S1A**), and we demonstrate here that *btaI*1 is directly controlled by BtaR1 (**Fig. S1B**). However, we were unable to witness a change in *btaI*1 activation by C_8_-HSL addition in our heterologous system, suggesting that an unknown factor could be involved in the interaction of BtaR1 with its ligand. While we demonstrated that BtaR1 constitutes the main regulator of *btaI*1 expression, we assume that BtaR1 represents the main regulator of C_8_-HSL biosynthesis as well. Since the Δ*btaR*1 mutant surprisingly accumulated more C_8_-HSL (**Fig. 2A**), posttranscriptional regulation on *btaI*1 expression is logically occurring. We propose that overproduction of C_8_-HSL detected in the Δ*btaR*1 mutant might be linked to RsaM1 activity. However, it is not excluded that additional unknown factors act on C_8_-HSL biosynthesis. A hypothetical protein encoded by the *BCAM1871* gene co-transcribed with *cepI*, contributes to QS in *B. cenocepacia* K56-2 (32). The BCAM1871 protein seems to act as an enhancer of AHL activity (32). Orthologs of the *BCAM1871* gene are found downstream from a gene encoding a LuxI-type synthase in many *Burkholderia* species (32). The *BTH_II1513* gene, in *B. thailandensis* E264, encodes a hypothetical protein sharing 56% identity with the BCAM1871 protein of *B. cenocepacia* J2315 (**Fig. S3**). Majerczyk *et al.* reported that *BTH_II1513* is activated by AHLs (19). Accordingly, our transcriptomic sequencing analyses confirm that QS regulates positively the *BTH_II1513* gene expression (S. Le Guillouzer, M. *C.* Groleau, and E. Déziel, unpublished data), and suggest that it is co-transcribed with *btaI*1. We propose that the BTH_II1513 protein is functionally homologous to the BCAM1871 protein, and could then similarly affect the QS-1 system.

The Δ*btaR*2 mutant also accumulated C_8_-HSL (**Fig. 2A**). Accordingly, we saw an increase in *btaI*1 expression in the absence of BtaR2 (**Fig. 2B**), which indicates again an interaction between QS-2 and QS-1 (**Fig. 8**), and reveals that the timing of expression of QS-1 system is dependent on QS-2. Our results also show that *btaI*1 expression was activated by 3OHC_10_-HSL and 3OHC_8_-HSL (**Fig. 6A**). Since no overexpression of the *btaI*1 gene was noticed in the Δ*btaI*2 mutant (**Fig. 2B**), we propose that BtaR2 represses the QS-1 in absence of its ligand (**Fig. 8**), suggesting that *btaI*1 activation by 3OHC_10_-HSL and 3OHC_8_-HSL involves alternative LuxR-type transcriptional regulators.

We have also shown that *btaI*1 is positively controlled by BtaR3 (**Fig. 2B**). Thus, BtaR3 could be responsible for activation of *btaI*1 by 3OHC_10_-HSL and 3OHC_8_-HSL. Consistently, adding exogenously 3OHC_10_-HSL or 3OHC_8_-HSL to the culture of the Δ*btaR*3 mutant did not affect expression of *btaI*1 (data not shown), confirming that activation of *btaI*1 by 3OHC_10_-HSL and 3OHC_8_-HSL is linked to BtaR3, revealing another interaction, this one between QS-3 and QS-1 (**Fig. 8**). Our results thus suggest that the QS-1 system could be positively controlled by BtaR3 at the transcriptional level. However, we observed an overproduction of C_8_-HSL in the Δ*btaR*3 mutant when compared to the wild-type strain (**Fig. 2A**), suggesting another regulation layer that as yet to be investigated. Additionally, *btaR*1 expression was upregulated in the Δ*btaI*1Δ*btaI*2Δ*btaI*3 mutant (**Fig. 5A**), indicating that expression of *btaR*1 is negatively regulated by C_8_-HSL, 3OH_10_-HSL, and 3OHC_8_-HSL. Accordingly, expression of the *bpsR* gene encoding BpsR was also reported to be QS regulated in *B. pseudomallei* (30).

The QS-3 system, homologous to *B. pseudomallei* BpsI3/BpsR3 and *B. mallei* BmaI3/BmaR3, is believed to be composed of BtaI3 and BtaR3 considering the juxtaposition of *btaI*3 and *btaR*3 on the genome of *B. thailandensis* (**Fig. S3**). The BtaI3 protein exhibits 92% identity with *B. pseudomallei* K96243 BpsI3 and *B. mallei* ATCC 23344 BmaI3, whereas the BtaR3 protein is 96% identical to BpsR3 and BmaR3. Similarly to the BpsI3 and BmaI3 synthases, BtaI3 was shown to produce 3OHC_8_-HSL (12, 13, 16). While the BpsR3 and BmaR3 transcriptional regulators specifically respond to 3OHC_8_-HSL, the *bpsI*3 and the *bmaI*3 genes were not reported to be directly activated by BpsR3 and BmaR3 in conjunction with 3OHC_8_-HSL, respectively (13, 16). Here in *B. thailandensis* E264, we demonstrated that *btaI*3 is positively controlled by BtaR3 and activated by 3OHC_8_-HSL since *btaI*3 expression is downregulated in the Δ*btaR*3 and Δ*btaI*3 mutants, respectively (**Fig. 4B**). Unexpectedly, transcription of *btaI*3 was restored in Δ*btaI*1Δ*btaI*2Δ*btaI*3 in the presence of 3OHC_8_-HSL, whereas adding 3OHC_8_-HSL to the cultures of the Δ*btaI*3 mutant had no impact on the *btaI*3 gene (**Fig. S2A**). Gamage *et al.* (16) demonstrated that the *B. pseudomallei* KHW BpsI/BpsR and BpsI3/BpsR3 QS circuits are both involved in the regulation of biofilm formation, whereas the development of biofilm was shown to be controlled by the QS-1 system in *B. thailandensis* E264 (33). While biofilm formation was reduced in the absence of the BpsI and BpsI3 synthases, and restored to wild-type levels in the presence of C_8_-HSL added exogenously to the culture of the *bpsI-* mutant strain, 3OHC_8_-HSL had not impact on the reduction in biofilm formed by the *bpsI3-* mutant (16). Interestingly, 3OHC_8_-HSL had also no impact on *btaI*3 in the Δ*btaI*1Δ*btaI*2Δ*btaI*3 mutant in the presence of C_8_-HSL and 3OHC_10_-HSL (**Fig. S5**). Collectively, these observations indicate that the QS-3 system is more complex than it appears. Considering that 3OHC_8_-HSL is not completely abolished in the absence of BtaI3, we hypothesize that the difference observed in *btaI*3 transcription between the *B. thailandensis* E264 wild-type strain and its Δ*btaI*3 mutant is not exclusively induced by 3OHC_8_-HSL, involving additional AHLs and/or alternative LuxR-type transcriptional regulators. Appropriately, we observed that *btaI*3 expression is also activated by 3OHC_10_-HSL, albeit to a lesser extent. Similarly, the BtaR3-controlled genes identified in transcriptomic analyses were also generally affected by both 3OHC_8_-HSL and 3OHC_10_-HSL (19). Considering that BpsR3 specifically responds to 3OHC_10_-HSL (16), and that BpsI3 and BmaI3 both produce 3OHC_10_-HSL in addition to 3OHC_8_-HSL (13, 16), it seems that BtaR3 also functions with both 3OHC_8_-HSL and 3OHC_10_-HSL. Additionally, *btaI*3 expression was substantially enhanced in the presence of 3OHC_8_-HSL (**Fig. 6C**), suggesting that, in contrast with BtaR2, the affinity of BtaR3 for 3OHC_8_-HSL is higher than for 3OHC_10_-HSL. Using the heterologous expression system *E. coli* DH5α, we also confirmed that BtaR3 directly activates *btaI*3 (**Fig. S2B**). Nevertheless, adding 3OHC_8_-HSL had no influence on *btaI*3 transcription. As suggested previously, the concentrations of exogenous 3OHC_8_-HSL added and/or the time exposure to this AHL chosen could be responsible (16). Another explanation could be disequilibrium between the artificially amounts of the BtaR3 protein produced and the levels of 3OHC_8_-HSL used in our experiments. We did not see any effect in the presence of 3OHC_10_-HSL as well (data not shown).

Interestingly, while activation of *btaI*3 by 3OHC_8_-HSL and 3OHC_10_-HSL occurred during the stationary phase (**Fig. 7**), auto-regulation of the *btaI*1 and *btaI*2 genes started during logarithmic growth. These results illustrate the sequential activation of the three QS systems observed during bacterial growth as QS-2 then QS-1 were both consecutively activated during the exponential phase, while *btaI*3 was not expressed until the stationary phase was reached (**Fig. 1B**). We thus hypothesize that the QS-3 system surrogates the regulation of the QS-2 system targets by producing 3OHC_8_-HSL in the stationary phase, whereas production of 3OHC_8_-HSL by the QS-2 system essentially occurs during the exponential phase, which would also explain why there is an overlap between both QS circuits when it comes to 3OHC_8_-HSL and 3OHC_10_-HSL modulated genes (19). Importantly, while sharing common AHLs, the QS-2 and QS-3 systems are apparently not transcriptionally linked, since BtaR2 does not regulate expression of *btaI*3, and the *btaI*2 gene is not controlled by BtaR3.

Since BtaR1 responds strongly to C_8_-HSL and that *btaI*3 is positively controlled by BtaR1, we concluded that BtaR1 regulates *btaI*3 expression in conjunction with C_8_-HSL. Furthermore, *btaI*3 was activated by C_8_-HSL during the exponential phase (**Fig. 7**), which is consistent with the idea that the QS-1 system is required for the expression of *btaI*3, and thus resulting in a belated activation of the QS-3 circuit and pointing toward an interaction between QS-1 and QS-3 (**Fig. 8**).

Additionally, the 3OHC_8_-HSL concentrations were increased in the Δ*btaR*1 mutant whereas expression of *btaI*3 was reduced to almost background levels. Consistently, expression of the *bpsI*3 gene, encoding the BpsI3 synthase, was also shown to be affected by the BpsI/BpsR QS system (16). Thus, these results indicate that the regulation of 3OHC_8_-HSL biosynthesis by BtaR1 does not implicate a direct interaction with the *btaI*3 promoter but rather could imply the effect of BtaR3 levels on *btaR*1 (19). We confirmed that, as seen for *btaI*3, *btaR*3 expression is activated by C_8_-HSL. These results support that *btaI*3 is indirectly controlled by BtaR1, highlighting again an interaction between QS-1 and QS-3 (**Fig. 8**). Furthermore, the *btaR*3 gene was also activated by 3OHC_8_-HSL and 3OHC_10_-HSL (**Fig. 5C**), suggesting that *btaR*3 expression is also auto-regulated.

## Conclusion

The study described here finally provides a more comprehensive and clearer picture of the interplay between the QS-1, QS-2, and QS-3 systems in *B. thailandensis* E264 (**Fig. 8**). We observed interdependence between the QS-1 and QS-2 systems that could involve additional factors such as the RsaM2 protein, highlighting once again that QS regulation can have many layers. While the QS-3 system was shown to be controlled by BtaR1, we also found that BtaR3 modulates the QS-1 system which indicates that those two systems are intertwined. Interestingly, such QS-1 and QS-3 systems interaction seems to be conserved in the closely-related species of the *Bptm* group (13, 16, 19). Additionally, the RsaM1 protein could play a role in the interconnection between the QS-1 and QS-3 systems. Interestingly, the QS-2 and QS-3 systems that share common AHLs are apparently not transcriptionally linked, but instead are temporally connected by their common AHLs. Collectively, our study suggests that there are homeostatic regulatory loops provided by the various QS systems in *B. thailandensis* resulting from transcriptional and post-transcriptional interactions, allowing tightly controlled coordination of gene expression.

Although we have found new connections and insights on the QS cascade, there are still many questions to be answered. The temporal pattern of QS-controlled genes clearly shows that additional factors are involved (19, 31, 34).

## Acknowledgments

Thank you to Everett Peter Greenberg (Department of Microbiology, University of Washington School of Medecine, Seattle, WA, USA) for providing *B. thailandensis* E264 strains. Special thanks to Sylvain Milot for his technical help.

## Legends for supplemental material

**Figure S1. *btaI*1 activation requires BtaR1 and C_8_-HSL.** (A) The luminescence of the mini-CTX-*btaI*1-*lux* transcriptional fusion was monitored during the exponential phase (OD_600_ ≈ 4.0) in cultures of the *B. thailandensis* E264 wild-type and the Δ*btaR*1, Δ*btaI*1, and Δ*btaI*1Δ*btaI*2Δ*btaI*3 mutant strains, haboring a *btaI*1-*lux* chromosomal reporter. Cultures were supplemented with 10 μM C_8_-HSL. Acetonitrile only was added in controls. The values represent the mean of three replicates. The luminescence is expressed in relative light units per culture optical density (RLU/OD_600_). (B) The luciferase activity of *btaI*1-*lux* was measured in the heterologous system *E. coli* DH5α, containing a BtaR1 expression vector with an arabinose-inducible promoter.

**Figure S2. *btaI*3 is activated by BtaR3 and 3OHC_8_-HSL.** (A) The luciferase activity of the mini-CTX-*btaI*3-*lux* transcriptional fusion was measured during the stationary phase (OD_600_ ≈ 8.0) in cultures of the *B. thailandensis* E264 wild-type and the Δ*btaR*3, Δ*btaI*3, and Δ*btaI*1Δ*btaI*2Δ*btaI*3 mutant strains, containing a *btaI*3-*lux* chromosomal reporter. Cultures were supplemented with 10 μM 3OHC_8_-HSL. Acetonitrile only was added in controls. The values represent the mean of three replicates. The luciferase activity is expressed in relative light units per culture optical density (RLU/OD_600_). (B) The luminescence of *btaI*3-*lux* was quantified in *E. coli* DH5α, expressing *btaR*3 controlled by an arabinose-inducible promoter.

**Figure S3. Genetic organization of the QS-regulatory genes in *B. thailandensis* E264.** *btaR*1 and *btaI*1 are not located next to each other and are divergently transcribed in *B. thailandensis* E264. The promoter region of *btaI*1 contains a putative *lux*-box sequence centered 73.5 bp usptream of the *btaI*1 translation start site (CCCTGTAAGGGTTAACAGTT). *btaR*2 and *btaI*2 are also not located next to each other and are divergently transcribed in *B. thailandensis* E264 as well. The promoter region of *btaI*2 contains a putative *lux*-box sequence centered 65.0 bp usptream of the *btaI*2 translation start site (ACCTGTAGAAATCGTAGT). *btaI*3 and *btaR*3 are transcribed in the same direction and are located next to each other in *B. thailandensis* E264.

**Figure S4. *btaI*2 is directly activated by BtaR2 in response to 3OHC_8_-HSL or 3OHC_10_-HSL.** The luciferase activity of the mini-CTX-*btaI*2-*lux* transcriptional fusion was monitored in the heterologous system *E. coli* DH5α, containing a BtaR2 expression vector with an arabinose-inducible promoter. Cultures were supplemented with 10 μM C_8_-HSL, 3OHC_8_-HSL, or 3OHC_10_-HSL. Acetonitrile only was added in controls. The values represent the mean of three replicates. The luminescence is expressed in relative light units per culture optical density (RLU/OD_600_).

**Figure S5. 3OHC_8_-HSL activation of *btaI*3 is dependent on C_8_-HSL and 3OHC_10_-HSL.** The luciferase activity of the mini-CTX-*btaI*3-*lux* transcriptional fusion was measured during the stationary phase (OD_600_ ≈ 8.0) in cultures of the *B. thailandensis* E264 wild-type and the Δ*btaI*1Δ*btaI*2Δ*btaI*3 mutant strains, containing a *btaI*3-*lux* chromosomal reporter. Cultures were supplemented with 10 μM C_8_-HSL, 3OHC_10_-HSL, and 3OHC_8_-HSL. Acetonitrile only was added in controls. The values represent the mean of three replicates. The luciferase activity is expressed in relative light units per culture optical density (RLU/OD_600_).

**Table S1. Bacterial strains used in this study.**

**Table S2. Plasmids used in this study.**

**Table S3. Primers used for PCR.**

**Table S4. Primers used for qRT-PCR.**

